# Effect of C6-HSL and chlorogenic acid on spoilage characteristics of *Serratia liquefaciens* in mutton

**DOI:** 10.64898/2026.01.20.700591

**Authors:** Yue Gu, Yueting Gu, Yixin Li, Jianjun Tian

## Abstract

Quorum sensing (QS) is a bacterial cell-cell communication system that coordinates group behaviors via self-produced signaling molecules. *Serratia liquefaciens*, a common mutton spoilage bacterium, precisely regulates its spoilage capacity through the AHL-mediated QS system. The results demonstrated that 20 μmol/mL C6-HSL concentration-dependently enhanced AHL activity 1.26-fold, increased biofilm formation by 51.55%, and elevated protease activity and siderophore production by 37% and 26.78%, respectively. In contrast, 80 μg/mL chlorogenic acid significantly inhibited AHL activity (by 26.19%), biofilm formation (by 42.54%), protease activity (by 28.92%), and motility (by 38.34%). In stored mutton, chlorogenic acid treatment reduced total plate counts by 6.1% and pH by 5.26%. Transcriptomic analysis revealed that C6-HSL treatment altered metabolic pathways such as flagellar assembly, ABC transporters, two-component systems, and secondary metabolite synthesis, in which spoilage-related genes (*slyB*, *fimA*, *fliJ*, *iucD*, *cheW*) were significantly up-regulated. In contrast, chlorogenic acid treatment affected pathways including amino acid metabolism, sulfur metabolism, and carbohydrate metabolism, with spoilage-related genes (*fimA*, *tuf*, *ibpA*, *clpS*, *metQ*) significantly down-regulated. These findings demonstrate that AHL activity plays a key role in regulating the spoilage capacity of *Serratia liquefaciens*, and suggest chlorogenic acid as a potential QS inhibitor with promising application in mutton preservation.

**Importance:** *Serratia liquefaciensis* a common spoilage bacterium in refrigerated foods and proliferates extensively in meat, making it one of the primary spoilage organisms. The significance of our research lies in investigating how changes in the activity of the signaling molecule AHL affect both spoilage capacity and mutton quality. Additionally, transcriptomic analysis was employed to elucidate the regulatory mechanisms by which altered AHL activity influences the spoilage potential of *S. liquefaciens*. The results demonstrated that variations in AHL activity significantly impacted key spoilage-related traits of *S. liquefaciens*, including biofilm formation, protease activity, and motility, while also contributing to improved meat quality during storage. Furthermore, the study revealed that AHL activity regulates metabolic pathways associated with spoilage as well as the expression of spoilage-related genes. These findings provide a theoretical basis for developing preservation strategies for mutton.

## Introduction

*Serratia liquefaciens* is a gram-negative bacillus and a common spoilage bacterium in refrigerated foods. It tends to proliferate extensively in mutton, becoming one of the predominant spoilage organisms. During its growth and metabolism, *S. liquefaciens* secretes hydrolytic enzymes that break down proteins and fats in the meat. This leads to sensory deterioration, including darkening of color and the development of off-odors, as well as nutrient loss (e.g., amino acids and vitamins), significantly reducing the edible quality and safety of mutton (Rood et al., 2022; Zhang et al., 2025).

Quorum sensing (QS) plays a crucial role in the spoilage process of mutton mediated by *S. liquefaciens* and other spoilage bacteria (Zaitseva et al., 2019). QS is a communication system wherein microbes produce and perceive diffusible signal molecules. This process allows the population to sense its density and accordingly orchestrate unified group behaviors (Tasneem et al., 2018; Han et al., 2022). In gram-negative bacteria, N-acyl homoserine lactones (AHLs) represent a typical class of QS signals (Papenfort & Bassler, 2016; Li et al., 2024). *S. liquefaciens* synthesizes and secretes multiple types of AHLs, and these AHL-mediated QS systems regulate various physiological traits—such as virulence factor expression, biofilm formation, and extracellular enzyme secretion—thereby enhancing its survival and spoilage potential in food matrices (Zheng et al., 2021; Sun et al., 2024). Therefore, investigating the activity of AHL signaling molecules in *S. liquefaciens* and their regulatory roles is essential for understanding the molecular mechanisms underlying microbial spoilage in mutton, as well as for developing novel meat preservation strategies that target QS (Patel et al., 2023; Fong et al., 2018).

The research aimed to investigate the effects of AHL signaling molecule activity changes on the spoilage characteristics of *Serratia liquefaciens* D1 and the storage quality of mutton. Integrated with transcriptomic analysis, the research revealed key genes and pathways through which the quorum sensing system regulates spoilage-related phenotypes at the molecular. The findings not only provide a solid foundation for elucidating the regulatory mechanism of the QS system in the spoilage traits of *S. liquefaciens*, but also offer important theoretical and practical references for developing mutton preservation strategies based on QS interference technology.

## Results and Discussion

### Effects of C6-HSL and chlorogenic acid on AHL activity and growth of *S. liquefaciens* D1

As shown in Fig. 1A, the effects of 20 μmol/mL C6-HSL and 80 μg/mL chlorogenic acid on the AHL activity of *S. liquefaciens* D1 were evaluated after 16 h of treatment. The results demonstrated that the addition of 20 μmol/mL exogenous C6-HSL significantly promoted AHL activity (*p* < 0.05), increasing it by 1.26-fold compared to the control —a finding consistent with that reported by Tang et al. (2015). In contrast, 80 μg/mL chlorogenic acid markedly inhibited AHL activity (*p* < 0.05), reducing it by 26.19%, which aligns with observations made by Zhang et al. (2020). However, neither compound significantly affected bacterial cell density (*p* > 0.05). These results indicate that C6-HSL and chlorogenic acid do not directly influence bacterial cells but instead function by modulating the levels of signaling molecules in the quorum sensing system.

**Fig. 1.**
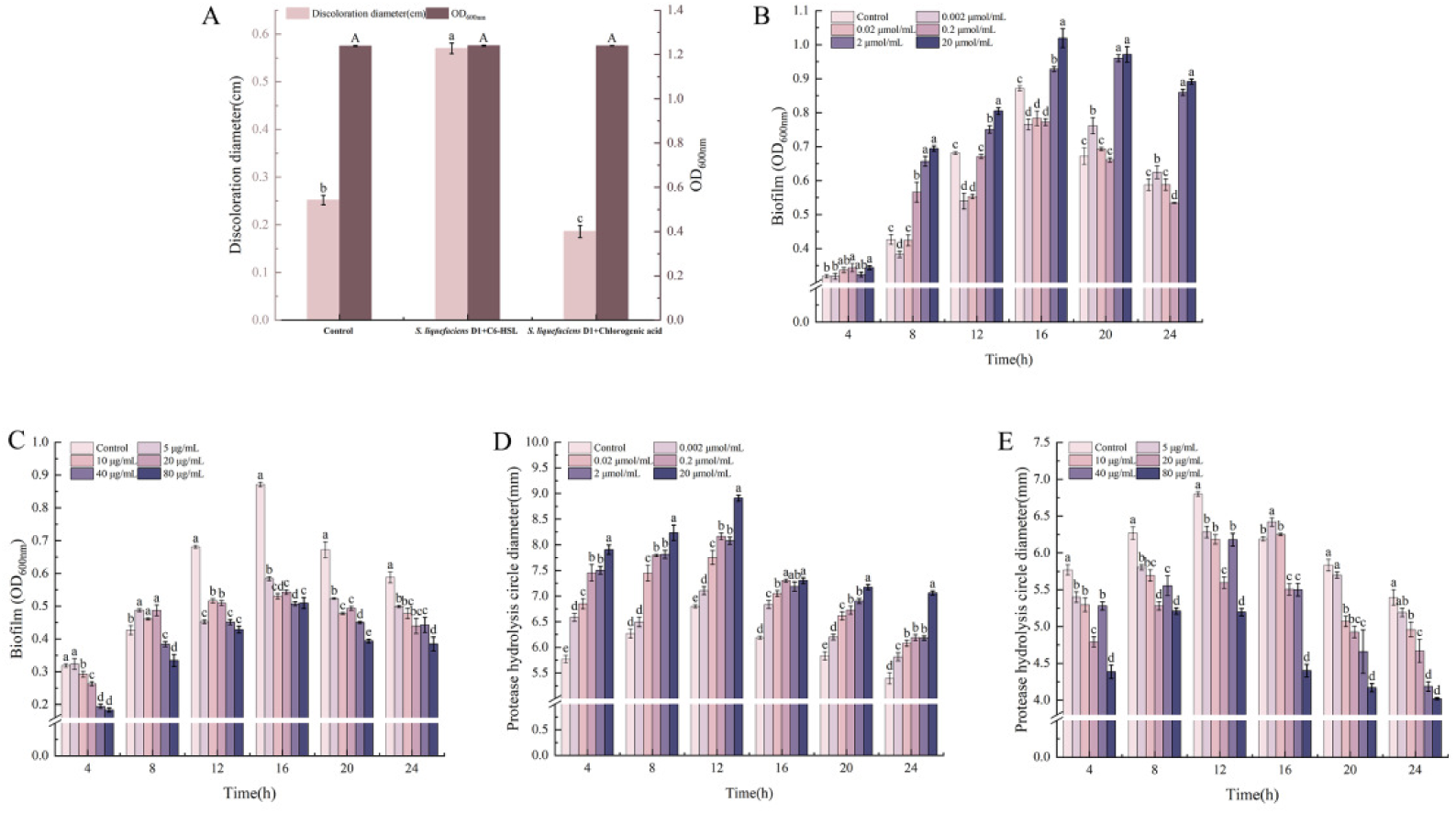
(A) Effects of C6-HSL and chlorogenic acid on AHL activity and cell density of *S. liquefaciens* D1. Effects of C6-HSL (B) and chlorogenic acid (C) on biofilm formation of *S. liquefaciens* D1. Effects of C6-HSL (D) and chlorogenic acid (E) on protease activity of *S. liquefaciens* D1. Data are presented as mean ± standard deviation. In Fig. 1A, uppercase and lowercase letters indicate significant differences between different treatment groups for the same measurement index (*p* < 0.05). In Figs. 1B, C, and D, different lowercase letters indicate significant differences between different concentrations at the same time point (*p* < 0.05).

### Effects of C6-HSL and chlorogenic acid on the spoilage phenotypes of *S. liquefaciens* D1

#### Effects of C6-HSL and chlorogenic acid on biofilm formation of *S. liquefaciens* D1

As shown in Fig. 1B, exogenous C6-HSL at concentrations of 0.002, 0.02, 0.2, 2, and 20 μmol/mL was applied to investigate its effects on the biofilm formation of *S. liquefaciens* D1. At 4 h, compared with the control, 0.2 and 20 μmol/mL C6-HSL significantly promoted biofilm formation (*p* < 0.05). At 8 h, 0.002 μmol/mL C6-HSL significantly inhibited biofilm formation, while all other concentrations exerted a significant promoting effect (*p* < 0.05). At 12 h, 0.002 and 0.02 μmol/mL C6-HSL significantly inhibited biofilm formation (*p* < 0.05), whereas 2 and 20 μmol/mL C6-HSL significantly enhanced it (*p* < 0.05). At 16 h, 0.002, 0.02, and 0.2 μmol/mL C6-HSL significantly suppressed biofilm formation (*p* < 0.05), while 2 and 20 μmol/mL C6-HSL showed a significant promoting effect (*p* < 0.05). At 20 h, 0.002, 2, and 20 μmol/mL C6-HSL significantly promoted biofilm formation (*p* < 0.05). Collectively, these results indicated that except for 4 h, 2 and 20 μmol/mL C6-HSL consistently promoted biofilm formation from 8 to 24 h. Specifically, biofilm production increased by 51.55% compared to the control at 24 h of incubation. These findings are consistent with those of Qian et al. (2021), who reported that the biofilm yield of *Pseudomonas aeruginosa* reached its maximum upon the addition of 25 µM C6-HSL. However, the inhibitory effect of low C6-HSL concentrations on biofilm formation observed in this study contradicts the results of Hou et al. (2017), who found that low concentrations of C6-HSL significantly promoted biofilm formation in *H. alvei* H4, whereas high concentrations slightly inhibited it. This discrepancy can be attributed to the fact that the effects of AHLs on biofilm formation vary among different bacterial species and are dependent on both dosage and type.

As shown in Fig. 1C, the effects of chlorogenic acid at concentrations of 5, 10, 20, 40, and 80 μg/mL on biofilm formation by *S. liquefaciens* D1 were evaluated within 24 h. At 4 h, chlorogenic acid concentrations ranging from 10 to 80 μg/mL significantly inhibited biofilm formation (*p* < 0.05). By 8 h, concentrations of 5–20 μg/mL significantly promoted biofilm formation (*p* < 0.05), while concentrations of 40–80 μg/mL significantly inhibited it (*p* < 0.05). From 12 to 24 h, all tested concentrations significantly inhibited biofilm formation (*p* < 0.05). The most pronounced inhibition was observed at 80 μg/mL (*p* < 0.05), which reduced biofilm formation by 42.54% at 4 h. This finding aligns with Wang et al. (2023), who reported that chlorogenic acid significantly inhibits biofilm formation in *P. putida* 433 at concentrations above 0.53 mM. The observed promotion of biofilm formation following 8 h treatment with 5–20 μg/mL chlorogenic acid may be attributed to a low-level stress response, whereby sub-inhibitory concentrations transiently stimulate adaptive bacterial mechanisms.

#### Effects of C6-HSL and chlorogenic acid on protease activity of *S. liquefaciens* D1

The protease activity (Fig. 1D) in both the control and various C6-HSL treatments showed an initial increase followed by a decline, peaking around 12 h. The effect of C6-HSL on protease activity exhibited concentration dependence. Except for the 0.002 μmol/mL C6-HSL treatment at 8 h, which showed no significant increase in protease activity compared to the control (*p* > 0.05), likely because the low concentration did not reach the signaling threshold and the bacterial density was insufficient to significantly induce protease activity, all other concentrations of C6-HSL significantly promoted protease activity in *S. liquefaciens* D1 at the remaining time points (*p* < 0.05). The strongest promotion was observed at a concentration of 20 μmol/mL, which increased protease activity by 37% compared to the control after 4 h of incubation. This finding is consistent with the research by Dai et al. (2022), who reported that the addition of 10 μmol/L C6-HSL significantly promoted protease activity in *Pseudomonas koreensis* PS1.

As shown in Fig. 1E, from 4 to 12 h, all tested concentrations of chlorogenic acid significantly inhibited protease activity in *S. liquefaciens* D1 (*p* < 0.05), with the inhibition exhibiting concentration dependence. At 16 h, 5 μg/mL chlorogenic acid promoted protease activity, while all other concentrations significantly inhibited it (*p* < 0.05). From 20 to 24 h, there was no significant difference in protease activity between the 5 μg/mL chlorogenic acid treatment and the control (*p* > 0.05), whereas all other concentration groups maintained significant inhibition of protease activity (*p* < 0.05). Among these, the 80 μg/mL chlorogenic acid treatment demonstrated the most significant inhibition throughout the entire process (*p* < 0.05), achieving a 28.92% inhibition rate on protease activity at 16 h of incubation. This result aligns with the findings of Wang et al. (2022), who observed that after adding 1/2 and 1/4 MIC of chlorogenic acid, the hydrolysis zone of the strain significantly decreased (*p* < 0.05), indicating that chlorogenic acid exerts an inhibitory effect on protease production. This may be related to the inhibition of signal molecule AHL activity by chlorogenic acid, thereby suppressing the spoilage characteristics associated with the strain.

#### Effects of C6-HSL and chlorogenic acid on motility and siderophore production of *S. liquefaciens* D1

As shown in Fig. 2A, as the concentration of C6-HSL increased, the motility of *S. liquefaciens* D1 gradually enhanced, with the most significant promotion (*p* < 0.05) observed at 20 μmol/mL, resulting in a 26.69% improvement in motility. Similarly, siderophore production also increased progressively with higher C6-HSL concentrations, reaching its most significant enhancement (*p* < 0.05) at 20 μmol/mL, with siderophore yield rising by 26.78%.

**Fig. 2.**
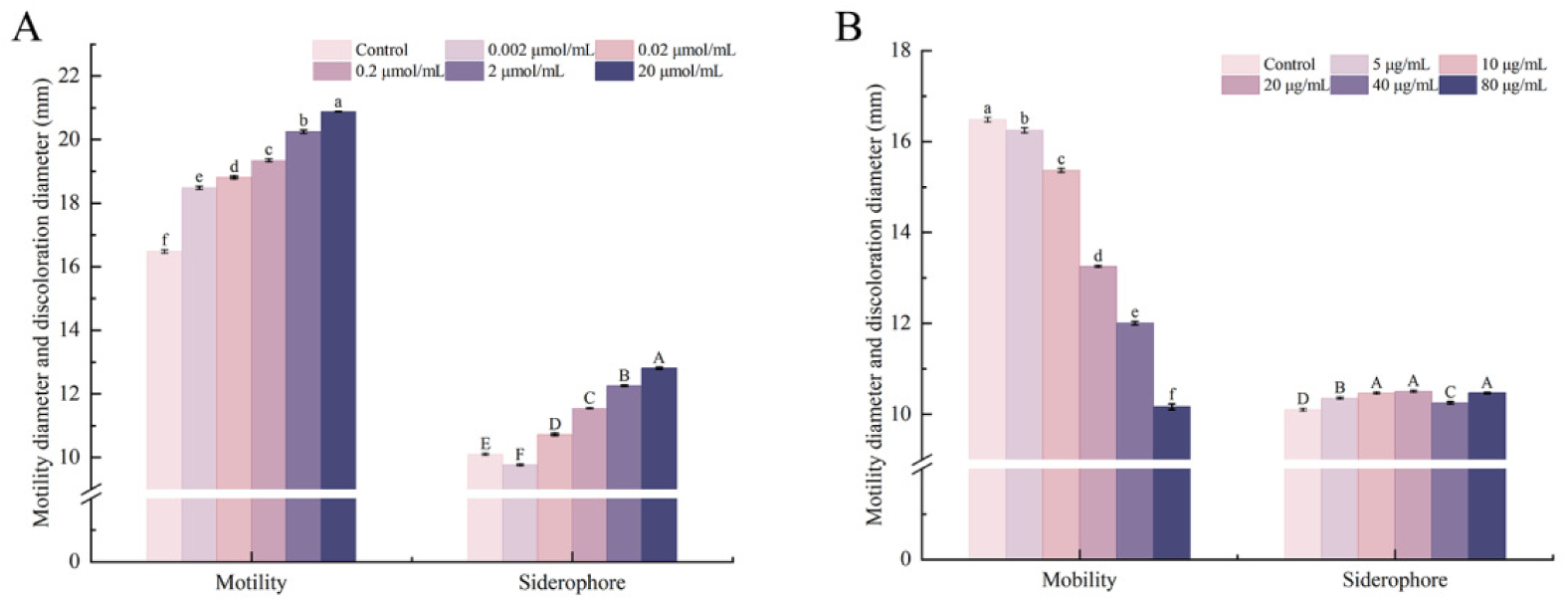
Effects of C6-HSL (A) and chlorogenic acid (B) on motility and siderophore production of *S. liquefaciens* D1. Different uppercase and lowercase letters indicate significant differences between different concentrations (*p* < 0.05).

As shown in Fig. 2B, increasing concentrations of chlorogenic acid led to a gradual reduction in motility, with the most significant inhibition (*p* < 0.05) observed at 80 μg/mL, resulting in an inhibition rate of 38.34%. In contrast, chlorogenic acid treatment showed no significant effect on siderophore production (*p* > 0.05). However, this finding contrasts with the results reported by Wang et al. (2020), who observed that chlorogenic acid treatment effectively suppressed siderophore production in *Pseudomonas fluorescens* in a concentration-dependent manner.

### Effects of chlorogenic acid on the quality of mutton during storage

#### Effects of chlorogenic acid on total plate count of mutton during storage

As shown in Fig. 3A, the effect of 80 μg/mL chlorogenic acid on the total plate count of mutton inoculated with *S. liquefaciens* D1 during storage was illustrated. With prolonged storage time, the total plate counts in both the control and chlorogenic acid-treated groups showed a gradual increasing trend. From days 3 to 9, the total bacterial count in the chlorogenic acid-treated group was significantly lower than that in the control group (*p* < 0.05). By day 9, the count in the treated group was 6.10% lower than that in the control, indicating that chlorogenic acid effectively inhibited bacterial growth during this stage and extended the shelf life of mutton. This finding is consistent with that of Liu et al. (2025), who reported that chlorogenic acid treatment significantly delayed the increase in total bacterial count in pork during storage, confirming its antibacterial effect. On day 12 of storage, there was no significant difference between the chlorogenic acid-treated group and the control (*p* > 0.05), suggesting that the antibacterial effect of chlorogenic acid had largely diminished by this point, with bacterial counts reaching similar levels. By day 15 of storage, the total plate count in the chlorogenic acid-treated group was significantly higher than that in the control, exceeding it by 3.74% (*p* < 0.05). This may be attributed to the potential degradation and utilization of chlorogenic acid by microorganisms or shifts in microbial community structure, which could have promoted the growth of specific microorganisms and led to a higher bacterial count in the treated group compared to the control in the later storage period.

**Fig. 3.**
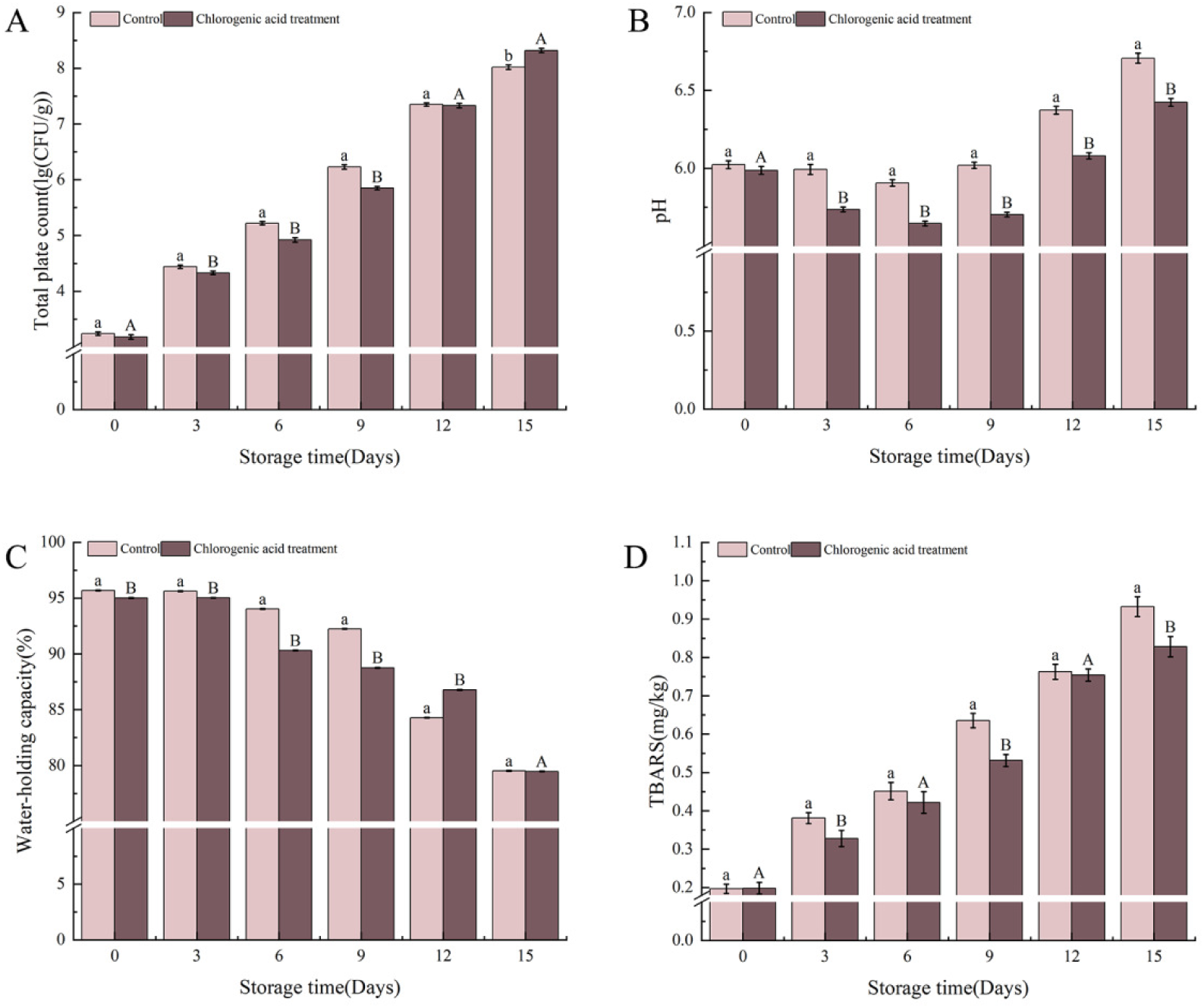
Effects of chlorogenic acid on total plate count (A), pH (B), water-holding capacity (C) and TBARS (D) of mutton during storage after inoculation with *S. liquefaciens* D1. Different uppercase and lowercase letters indicate significant differences between treatment and the control at the same time point (*p* < 0.05).

#### Effects of chlorogenic acid on pH of mutton during storage

The pH levels in both the chlorogenic acid-treated and the control (Fig. 3B) exhibited an initial decline followed by a gradual increase over time, ranging overall between 5.6 and 6.7. During the early storage period (0–6 days), the decrease in pH in both groups was primarily attributed to the production of lactic acid and other organic acids from glycogen glycolysis under near-anaerobic conditions. On day 6, the pH of both the control and treated groups reached their minimum values, measuring 5.90 and 5.65, respectively. In the mid-to-late storage stage (6–15 days), the pH trend reversed and began to rise, reaching peak values of 6.7 and 6.42 for the control and treated groups, respectively, on day 15. This increase can be attributed to the synergistic action of microorganisms, endogenous and exogenous enzymes accelerating protein degradation, as well as the ongoing catabolic processes that consumed acidic metabolites such as lactic acid. From days 3 to 15, the pH values of the chlorogenic acid-treated group remained significantly lower than those of the control (*p* < 0.05), and were 5.26% lower than the control on day 12, indicating that chlorogenic acid effectively delayed the pH increase during mutton storage. However, a study by Zhu et al. (2024) found that chlorogenic acid treatment had no significant effect on the pH of tuna meat.

#### Effect of chlorogenic acid on water-holding capacity of mutton during storage

With the extension of storage time, the water-holding capacity of both the control and the treatment group (Fig. 3C) showed a declining trend. During the storage period of 0–9 days, the water-holding capacity of the chlorogenic acid-treated group was significantly lower than that of the control (*p* < 0.05). By day 9, the treated group was 3.73% lower than the control. This may be attributed to chlorogenic acid as a phenolic acid, whose slight acidity or interaction with muscle proteins could have initially affected the charge and hydration state of myofibrillar proteins, leading to a slightly higher release of free water compared to the control. On day 12 of storage, the water-holding capacity of the chlorogenic acid-treated group was significantly higher than that of the control (*p* < 0.05), exceeding the control by 2.92%. This may be because chlorogenic acid can delay the oxidation of fats and proteins, inhibit microbial growth, and thereby maintain the integrity of muscle cell structure and the water-retaining capacity of proteins. By day 15 of storage, there was no significant difference between the two groups (*p* > 0.05), indicating that with further extension of storage time, protein oxidation and damage to muscle tissue structure had reached a high level, and the preservative effect of chlorogenic acid was gradually offset. However, Liu et al. (2025) found that under non-drained storage conditions, chlorogenic acid treatment did not result in a significant difference in water loss rate compared to the control in pork.

#### Effect of chlorogenic acid on TBARS of mutton during storage

With the extension of storage time, the TBARS values of both the control and treatment group (Fig. 3D) showed an increasing trend. On days 3, 9, and 15 of storage, the TBARS values in the chlorogenic acid-treated group were significantly lower than those in the control (*p* < 0.05), being 13.91%, 16.37%, and 11.16% lower, respectively. This indicates that chlorogenic acid can inhibit lipid oxidation to some extent and delay the quality deterioration of mutton. On days 6 and 12 of storage, there was no significant difference between the chlorogenic acid-treated group and the control (*p* > 0.05). This could be attributed to nonlinear changes in the lipid oxidation process or a temporary equilibrium reached between the antioxidant efficacy of chlorogenic acid and environmental factors at these time points. Xie et al. (2021) found that chlorogenic acid could inhibit lipid oxidation in precooked white shrimp, demonstrating a certain preservative effect.

#### Effect of chlorogenic acid on color and texture of mutton during storage

As shown in Table 1, during the storage of mutton, the brightness (L*) and redness (a*) values of both the control and the chlorogenic acid-treated group exhibited a decreasing trend over time, although the decline was alleviated in the treated group. This indicates that chlorogenic acid slowed the surface darkening of the meat samples. The L* value of the control decreased from an initial 48.83 to 35.81, while that of the treated group decreased from 49.36 to 36.24. On day 6 of storage, the L* value of the chlorogenic acid-treated group was significantly higher than that of the control, exceeding it by 14.04% (*p* < 0.05), while no significant differences were observed at other time points (*p* > 0.05). The a* value of the control decreased from an initial 18.61 to 6.17, while that of the treated group decreased from 18.39 to 7.05. Although the a* value of the treated group was overall higher and declined more gradually than that of the control, the difference was not statistically significant (*p* > 0.05). This may be because changes in the redness of mutton are influenced by multiple factors, including myoglobin oxidation, lipid peroxidation, and photolysis, and the singular antioxidant effect of chlorogenic acid may not fully dominate this complex process.

**Table 1.** Effect of chlorogenic acid on the color of mutton inoculated with *S. liquefaciens* D1 during storage.

|  |  | Storage time (Days) |  |  |  |  |  |
| --- | --- | --- | --- | --- | --- | --- | --- |
|  |  | 0 | 3 | 6 | 9 | 12 | 15 |
| L* | Control | 48.83±0.27<sup>a</sup> | 47.64±0.28<sup>a</sup> | 41.17±0.32<sup>a</sup> | 39.97±0.34<sup>a</sup> | 38.42±0.35<sup>a</sup> | 35.81±0.31<sup>a</sup> |
|  | Treatment | 49.36±0.36<sup>a</sup> | 48.52±0.29<sup>a</sup> | 46.95±0.29<sup>b</sup> | 40.77±0.25<sup>a</sup> | 40.16±0.28<sup>b</sup> | 36.24±0.27<sup>a</sup> |
| a* | Control | 18.61±0.42<sup>a</sup> | 16.94±0.33<sup>a</sup> | 14.98±0.21<sup>a</sup> | 8.39±0.21<sup>a</sup> | 6.96±0.18<sup>a</sup> | 6.17±0.29<sup>a</sup> |
|  | Treatment | 18.39±0.37<sup>a</sup> | 17.42±0.29<sup>a</sup> | 15.66±0.29<sup>a</sup> | 8.03±0.25<sup>a</sup> | 7.64±0.25<sup>a</sup> | 7.05±0.24<sup>a</sup> |

Note: L\* indicates lightness, and a\* indicates redness. Different letters indicate significant differences
( $p < 0.05$ ).

As shown in Table 2, the hardness, springiness, and chewiness of mutton samples in both the control and treatment groups decreased with prolonged storage time. Chlorogenic acid treatment partially slowed the decline in these textural properties. However, no statistically significant differences in texture parameters were observed between the groups throughout the storage period (*p* > 0.05), indicating that the treatment did not have a significant impact on the texture of the mutton. This may be because the antioxidant effect of chlorogenic acid did not effectively delay the denaturation and degradation of muscle proteins.

**Table 2.** Effect of chlorogenic acid on the texture of mutton during storage.

|  |  | Storage time (Days) |  |  |  |  |  |
| --- | --- | --- | --- | --- | --- | --- | --- |
|  |  | 0 | 3 | 6 | 9 | 12 | 15 |
| Hardness<br>(g) | Control | 143.56±14.25<sup>a</sup> | 114.62±17.21<sup>a</sup> | 83.16±20.18<sup>a</sup> | 72.22±23.42<sup>a</sup> | 63.18±19.34<sup>a</sup> | 52.22±19.23<sup>a</sup> |
|  | Treatment | 149.14±16.42<sup>a</sup> | 106.25±19.38<sup>a</sup> | 78.46±21.42<sup>a</sup> | 78.26±19.42<sup>a</sup> | 72.83±20.48<sup>a</sup> | 60.15±18.29<sup>a</sup> |
| Elasticity<br>(mm) | Control | 0.91±0.02<sup>a</sup> | 0.88±0.03<sup>a</sup> | 0.72±0.06<sup>a</sup> | 0.68±0.03<sup>a</sup> | 0.52±0.03<sup>a</sup> | 0.42±0.04<sup>a</sup> |
|  | Treatment | 0.95±0.05<sup>a</sup> | 0.89±0.05<sup>a</sup> | 0.79±0.04<sup>a</sup> | 0.75±0.04<sup>a</sup> | 0.63±0.04<sup>a</sup> | 0.53±0.06<sup>a</sup> |
| Chewiness<br>(gmm) | Control | 122.61±9.31<sup>a</sup> | 92.65±5.19<sup>a</sup> | 67.77±5.48<sup>a</sup> | 46.32±7.26<sup>a</sup> | 42.32±6.54<sup>a</sup> | 39.98±7.22<sup>a</sup> |
|  | Treatment | 134.59±8.45<sup>a</sup> | 97.77±6.32<sup>a</sup> | 71.83±5.72<sup>a</sup> | 53.38±6.98<sup>a</sup> | 49.45±7.38<sup>a</sup> | 45.27±6.79<sup>a</sup> |

Note: Different letters within the same column indicate significant differences ( $p < 0.05$ ).

### Transcriptomic Analysis

#### Differential Gene Expression Analysis

The transcriptomes of *S. liquefaciens* D1 treated with C6-HSL and chlorogenic acid were compared to the untreated control, respectively. As a quorum-sensing signal molecule, C6-HSL significantly upregulated 584 genes and downregulated 25 genes (Fig. 4A). In contrast, chlorogenic acid, as a signal molecule inhibitor, resulted in the significant upregulation of 185 genes and downregulation of 119 genes (Fig. 4B). The cluster heatmaps (Fig. 4C and D) further confirmed that both treatments induced distinct gene expression profiles clearly separated from the control, validating the reliability of the treatment effects.

**Fig. 4.**
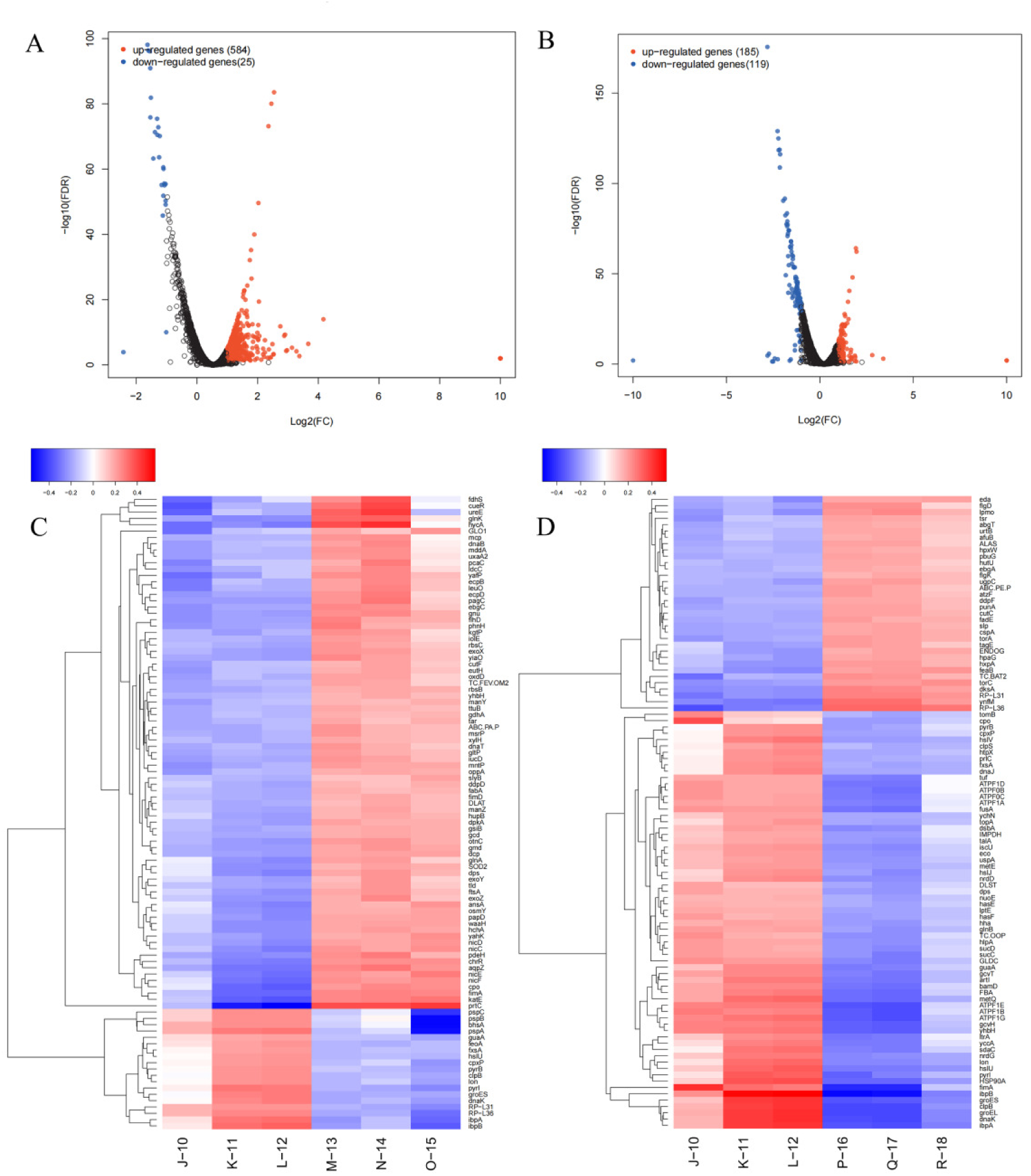
Analysis of differentially expressed genes (DEGs). (A) Volcano plot of all DEGs between the C6-HSL treatment and the control. (B) Volcano plot of all DEGs between the chlorogenic acid treatment and the control. (C) Cluster heatmap of DEGs between the C6-HSL treatment and the control. (D) Cluster heatmap of DEGs between the chlorogenic acid treatment and the control. (Control : J-10, K-11, L-12; C6-HSL treatment : M-13, N-14, O-15; Chlorogenic acid treatment : P-16, Q-17, R-18).

#### Functional annotation of differentially expressed genes

GO enrichment-based cluster analysis revealed significant enrichment in three main categories: biological process (BP), molecular function (MF), and cellular component (CC). Following C6-HSL treatment (Fig. 5A), differentially expressed BP genes were primarily associated with cellular process, metabolic process, response to stimulus, biological regulation, localization, multi-organism process, and interspecies interaction between organisms. For MF, the enriched categories included binding, catalytic activity, transporter activity, structural molecule activity, transcription regulator activity, protein folding chaperone, and antioxidant activity. In terms of CC, significant terms involved cellular anatomical entity, intracellular components, protein-containing complex, and other organism parts. Notably, C6-HSL treatment led to the enrichment of 11 BP, 4 MF, and 5 CC GO terms. In contrast, chlorogenic acid treatment (Fig. 5B) resulted in differentially expressed BP genes mainly involved in cellular process, metabolic process, response to stimulus, localization, biological regulation, multi-organism process, and growth. Enriched MF categories included binding, transporter activity, transcription regulator activity, molecular function regulator, structural molecule activity, protein folding chaperones, and translation regulator activity. For CC, the terms were related to cellular anatomical entity, intracellular structures, protein-containing complex, and other organism parts. This treatment led to a broader enrichment profile, with 49 BP, 29 MF, and 13 CC GO terms identified.These results indicate that both C6-HSL and chlorogenic acid predominantly influence biological processes in the tested bacterial strain.

**Fig. 5.**
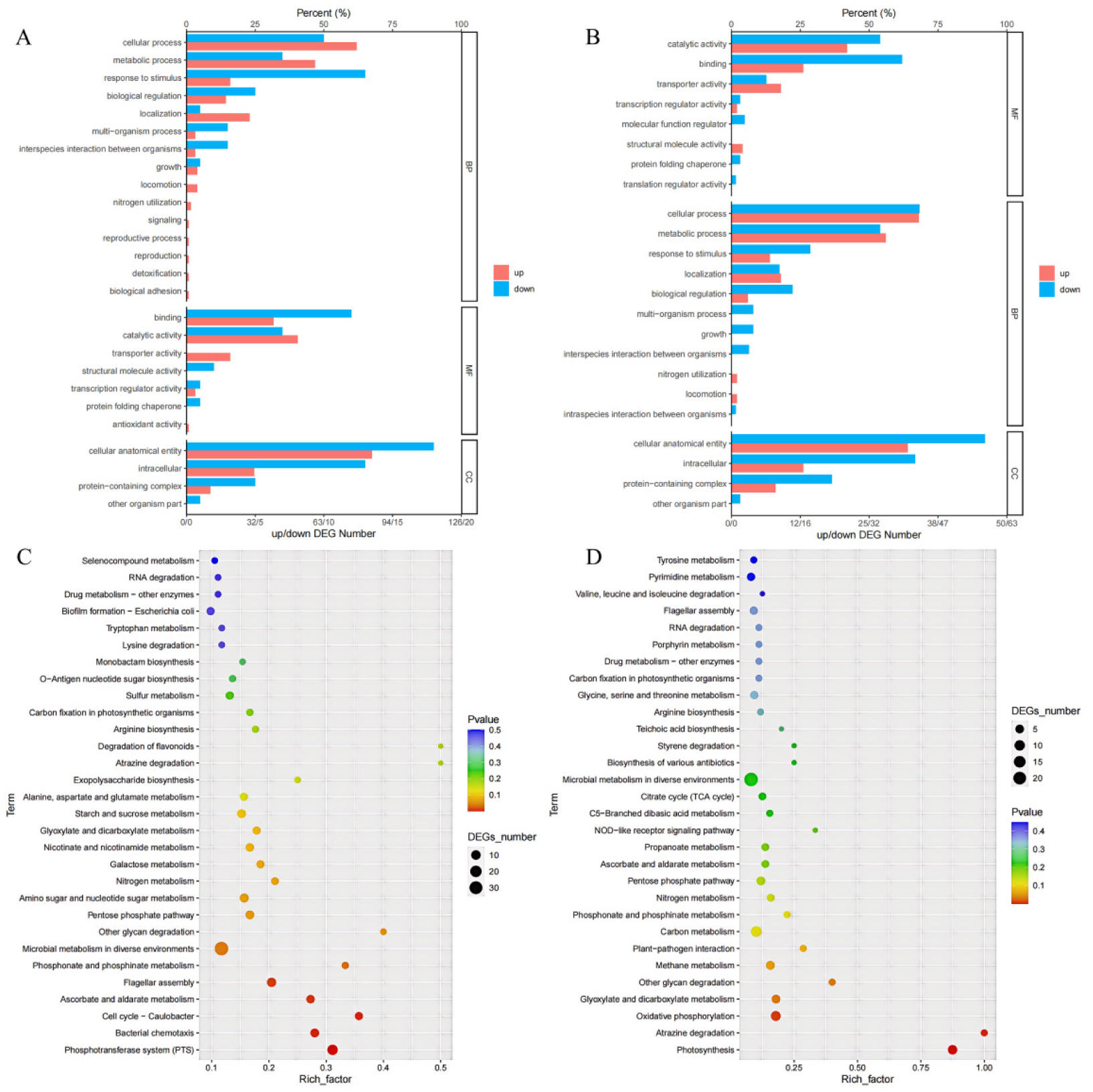
(A) Bar plot of GO enrichment results for the C6-HSL treatment versus the control. (B) Bar plot of GO enrichment results for the chlorogenic acid treatment versus the control. (C) Scatter plot of KEGG pathway enrichment analysis for differentially expressed genes (DEGs) in the C6-HSL treatment versus the control. (D) Scatter plot of KEGG pathway enrichment analysis for DEGs in the chlorogenic acid treatment versus the control. In panels C and D, the x-axis represents the rich factor, and the y-axis represents the KEGG pathways. The size of the bubble corresponds to the number of DEGs enriched in a given pathway.

Further KEGG pathway enrichment analysis was performed on the differentially expressed genes using a significance threshold of *p* < 0.05, and a scatter plot was generated for the top 30 most significant KEGG pathways. After C6-HSL treatment , a total of 328 differentially expressed genes were enriched in 75 KEGG pathways. In contrast, chlorogenic acid treatment resulted in 178 differentially expressed genes enriched in 66 KEGG pathways. C6-HSL significantly up-regulated genes associated with several key pathways (Fig. 5C), including the phosphotransferase system, flagellar assembly, microbial metabolism in diverse environments, ABC transporters, two-component system, and biosynthesis of secondary metabolites. Conversely, it significantly down-regulated genes related to cysteine and methionine metabolism, ribosome function, and pyrimidine metabolism. Similarly, chlorogenic acid significantly up-regulated genes involved in multiple metabolic processes (Fig. 5D), such as alanine, aspartate and glutamate metabolism, pentose and glucuronate interconversions, nitrogen metabolism, and sulfur metabolism. Meanwhile, it significantly down-regulated genes associated with oxidative phosphorylation, carbon metabolism, and carbohydrate metabolism.

### Analysis of key differentially expressed genes

Based on gene functional annotation information, functional genes related to spoilage characteristics and quorum sensing that showed differential expression in *S. liquefaciens* D1 after treatment with C6-HSL and chlorogenic acid were screened.

Differential gene expression analysis revealed that C6-HSL treatment (Table 3) significantly upregulated the expression of multiple genes associated with virulence and environmental adaptation. In terms of biofilm formation, the expression of genes such as *slyB*, *ecpB*, *ecpC*, *ecpD*, *fimA*, *fimD*, and *wzc* was enhanced. Among these, *slyB* contributes to maintaining biofilm integrity (Plesa et al., 2006), *wzc* is a known biofilm-promoting gene (Azra et al., 2025), and *fimA* has been confirmed as an important virulence factor in *Bacteroides nodosus* (Kennan et al., 2001). Flagellar synthesis-related genes (*fliZ*, *fliJ*, *fliQ*, *flhC*, *flhD*) were also significantly upregulated. As a key structure for bacterial motility and initial adhesion, flagella play a decisive role in biofilm formation (Li et al., 2018). Among these genes, Nguyen et al. (2016) found that *fliJ* is crucial for flagellar assembly, playing important roles not only in motility but also in virulence. Additionally, the expression of siderophore synthesis genes *iucD* and *fhuE* increased. *iucD* is involved in regulating iron uptake systems (Paixão et al., 2016). Furthermore, two-component system-related genes *cheW* and *tsr* were significantly upregulated. This system participates in regulating multiple key processes, including bacterial chemotaxis, biofilm formation, quorum sensing, and stress adaptation (Li et al., 2023). Notably, Shahbaz et al. (2020) found that deleting the *cheW1* gene in *Xanthomonas oryzae* pv resulted in reduced virulence.

**Table 3.**
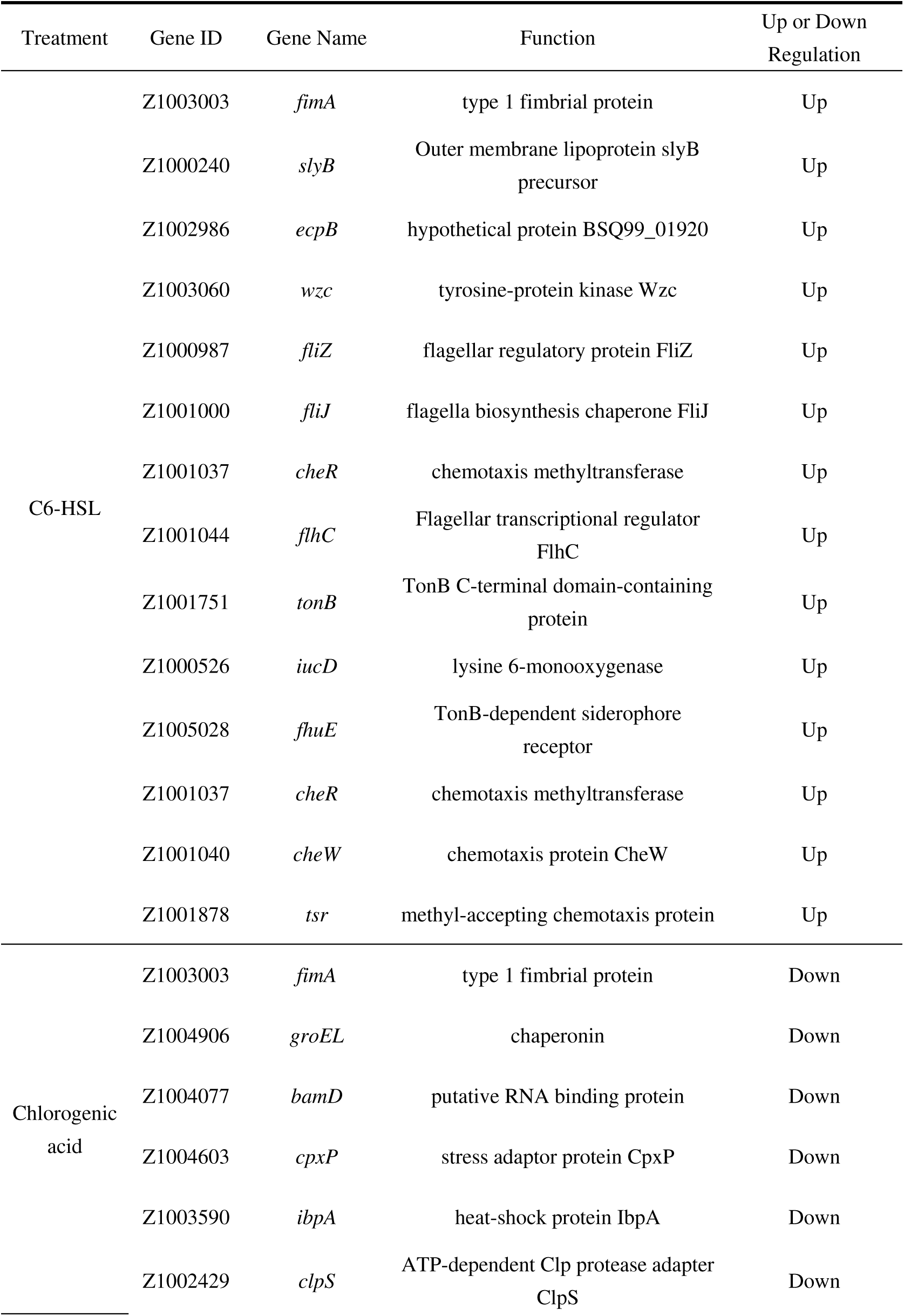
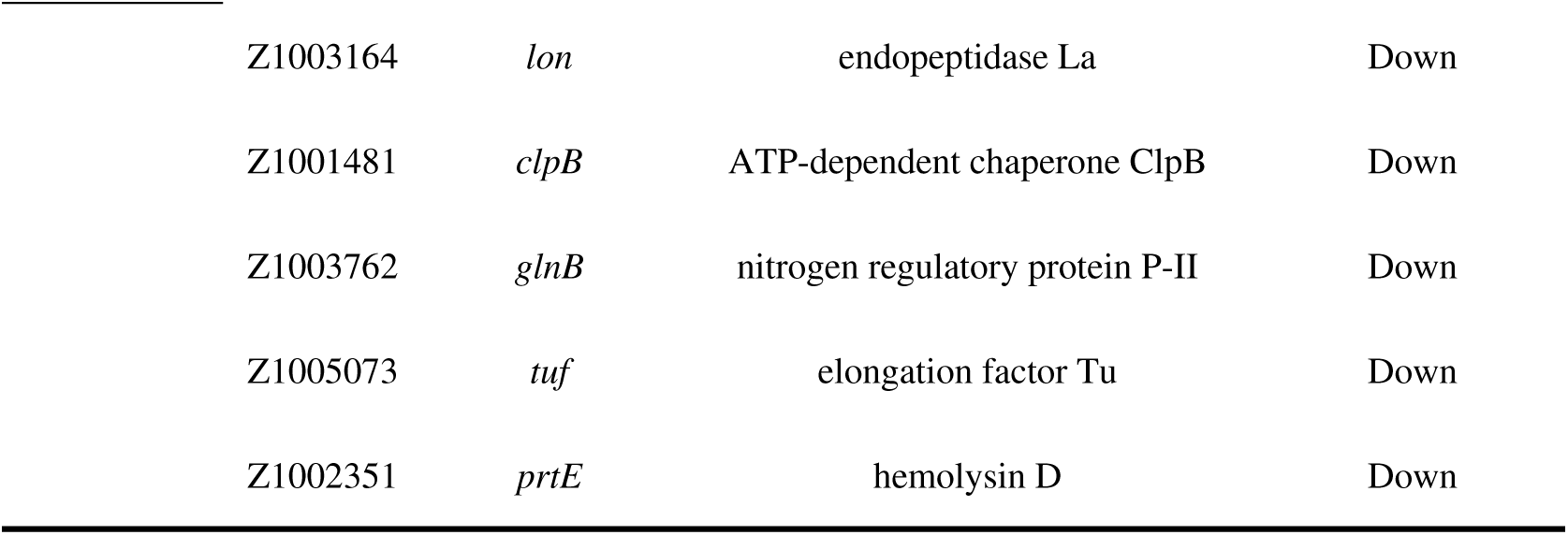
Significantly differentially expressed genes related to spoilage characteristics under C6-HSL or chlorogenic acid treatment.

After chlorogenic acid treatment (Table 3), the expression of several genes related to biofilm formation and stress response (such as *fimA*, *tuf*, *groEL*, *cpxP*, and *ibpA*) was down-regulated, indicating that bacterial adhesion capacity and environmental adaptability were suppressed. Among these, the *tuf* gene encodes the elongation factor Tu (EF-Tu). In *Gallibacterium*, EF-Tu may be involved in biofilm formation and is associated with bacterial pathogenicity (López-Ochoa et al., 2017). Additionally, Sun et al. (2020) found that knockout of the *ibpA* gene in *Escherichia coli* led to reduced biofilm formation. Furthermore, the expression levels of several protease system-related genes (such as *clpS*, *lon*, *clpB*, and *prtE*) also decreased, affecting intracellular protein homeostasis and the activity of enzymes related to spoilage processes. According to Lal et al. (2018), the ClpS caseinolytic protease encoded by *clpS* participates in the protein hydrolysis process. Additionally, genes related to the ABC transport system (*metQ* and *artI*) and the two-component system (*glnB*) showed significant down-regulation. Among these, Bosch et al. (2025) found that MetQ, encoded by *metQ*, plays a crucial role in methionine uptake, virulence, antioxidant stress response, and biofilm formation in *S. suis*.

## Conclusion

This study reveals the critical role of AHL signaling activity in regulating spoilage phenotypes and maintaining mutton quality during storage. The exogenous addition of the signaling molecule C6-HSL significantly up-regulated the expression of *S. liquefaciens* D1 genes related to biofilm formation, protease synthesis, flagellar assembly, and siderophore uptake. This enhanced the adhesion, motility, and nutrient acquisition efficiency of *S. liquefaciens* D1, ultimately increasing its spoilage potential. In contrast, chlorogenic acid, as a quorum sensing inhibitor, down-regulated the same categories of genes along with stress-response-related genes. This effectively suppressed the colonization ability and environmental adaptability of *S. liquefaciens* D1, thereby reducing its spoilage activity. These changes in gene expression levels further mediated spoilage-related phenotypes of the strain, such as protease activity, biofilm formation capacity, and motility, which were ultimately reflected in key quality indicators of mutton during storage, including pH changes and total plate count. From the perspective of signaling molecule regulation, this study systematically elucidates the microbial spoilage mechanism based on quorum sensing, providing a theoretical basis for the development of novel targeted preservation technologies, particularly those based on quorum sensing quenching. It offers certain theoretical value for achieving precise quality control during mutton storage.

## Material and Methods

### Strain and growth conditions

*S. liquefaciens* D1 was inoculated into Luria-Bertani (LB) broth and cultured aerobically in a shaking incubator at 33 ℃ with 150 rpm agitation for 16 h. This subculturing process was repeated twice to ensure culture purity and activate the bacterial growth.

### Effects of C6-HSL and chlorogenic acid on *S. liquefaciens* D1 growth

LB liquid medium was supplemented with 20 μmol/mL C6-HSL or 80 μg/mL chlorogenic acid, respectively. Bacterial strains from the second activation cycle were inoculated into the LB medium and cultured at 33 °C with shaking at 150 rpm for 16 h. The optical density of the bacterial culture at 600 nm (OD₆₀₀) was measured to assess growth.

### Effects of C6-HSL and chlorogenic acid on AHL activity in *S. liquefaciens* D1

*S. liquefaciens* D1 cultures were centrifuged (8000 × g, 5 min). The supernatant was filtered and extracted three times with ethyl acetate containing 0.01% glacial acetic acid. The organic phase was dried, and the residue was redissolved in 70% methanol to obtain crude AHL extract. For detection, *C. violaceum* CV026 agar plates were prepared, wells were punched, and 40 μL extract was added per well. After overnight incubation at 33 °C, violet halos were recorded. Reporter strain supernatant served as the negative control (Li et al., 2019).

### Effects of C6-HSL and chlorogenic acid on the spoilage phenotypes of *S. liquefaciens* D1

*S. liquefaciens* D1 was inoculated into LB broth supplemented with varying concentrations of either C6-HSL (0.002, 0.02, 0.2, 2, and 20 μmol/mL) or chlorogenic acid (5, 10, 20, 40, and 80 μg/mL). All cultures were then incubated at 33 °C with shaking at 150 r/min.

### Biofilm formation

Biofilm formation was quantified by crystal violet staining. Briefly, 160 µL LB medium, 20 µL of C6-HSL or chlorogenic acid (varying concentrations), and 20 µL of a 16 h bacterial culture were combined per well in a 96-well plate. After static incubation at 33 °C for 24 h, planktonic cells were removed, and wells were gently washed with PBS. Adherent cells were fixed with methanol, stained with crystal violet, and dissolved in glacial acetic acid. Absorbance at 595 nm (OD₅₉₅) was measured to quantify biofilm biomass (Mao et al., 2023).

### Protease activity

A 15% (w/v) skim milk solution was sterilized at 105 °C for 30 min, and a 1.5% (w/v) agar solution was autoclaved at 121 °C for 15 min. For the assay, 10 mL of the sterilized skim milk solution was mixed with 90 mL of molten agar. After the mixture solidified in plates, 5 μL of bacterial culture was spotted onto the agar surface, air-dried under sterile conditions, and incubated at 33 °C for 24 h. The diameter of the proteolytic halo was measured (Anbu et al., 2013).

### Motility and siderophore production

Swimming motility of *S. liquefaciens* D1 was assessed by spotting bacteria onto soft agar plates and incubating statically at 33 °C for 24 h; the diameter of the spreading zone was recorded (Jahid et al., 2015). For siderophore production, the strain was cultured overnight in skim milk medium at 33 °C with shaking. Subsequently, 60 µL of culture was applied to Chrome Azurol S (CAS) agar plates and incubated statically at 33 °C for 24 h; the diameter of the orange halo indicating siderophore secretion was measured (Shin et al., 2001).

### Effect of chlorogenic acid on mutton quality during storage

#### Meat sample preparation and bacterial inoculation

Fresh mutton was surface-sterilized with 75% ethanol, rinsed with sterile water, blotted dry, and UV-irradiated for 15 min on each side. The meat was aseptically cut into 5 g pieces, which were immersed for 1 min in sterile water (control) or 80 μg/mL chlorogenic acid solution. The pieces were then surface-inoculated with 10⁴ CFU/mL *S. liquefaciens* D1 suspension, air-dried for 30 seconds, vacuum-packaged, and stored at 4 °C (Zhao et al., 2025).

#### Total plate count

The meat sample (5 g) was homogenized with 45 mL of sterile saline solution. One milliliter of the homogenate was transferred into a test tube containing 9 mL of sterile physiological saline to perform serial dilutions. One milliliter of the diluted sample was then pipetted into a sterilized Petri dish, mixed with poured Plate Count Agar (PCA), and allowed to solidify. After solidification, the plates were incubated at 33 °C for 48 h for total plate count enumeration (Zuo et al., 2025).

#### Physicochemical indicators

Color measurement: Color difference was determined using a chroma meter (CR-400, Konica Minolta Sensing Americas, USA) and expressed as a* (redness) and L* (lightness) values (Ben Slama et al., 2017).

Texture analysis: A texture analyzer (TMS-Pro, Food Technology Corporation, USA) was used according to the method described by Spaziani et al. (2008).

Lipid oxidation: The extent of lipid oxidation was quantified using the thiobarbituric acid reactive substances (TBARS) method. Briefly, 5 g of meat was homogenized with 10 mL of 20% trichloroacetic acid (TCA), vortexed, and centrifuged at 3500 r/min for 15 minutes. The supernatant was filtered, and 5 mL was mixed with an equal volume of 0.02 mol/L TBA solution. The mixture was heated in a 90 °C water bath for 30–40 minutes, cooled, and the absorbance was measured at 532 nm (Chi et al., 2024).

pH measurement: pH was measured using a portable pH meter (FE28, Mettler Toledo, Switzerland). Ten grams of mutton was homogenized with 100 mL of distilled water prior to measurement (Yang et al., 2024).

Water-holding capacity (WHC): WHC was determined using a low-speed centrifugation method. Approximately 5 g of meat wrapped in double-layer qualitative filter paper was placed in a centrifuge tube and centrifuged at 8000 × g for 15 minutes. The mass of the meat sample was measured after centrifugation (Sánchez-Alonso et al., 2012).

#### Transcriptomic analysis

*S. liquefaciens* D1 was cultured and treated with either 20 μmol/mL C6-HSL or 80 μg/mL chlorogenic acid in LB medium at 33 °C for 16 h. Cells were harvested by centrifugation, washed twice with PBS, and flash-frozen in liquid nitrogen. PBS-treated samples served as the control. Each condition included three biological replicates.

Total RNA was extracted and assessed for purity (OD_260/280_ ratio of 1.8–2.2), integrity, and quantity. rRNA was depleted, and the enriched mRNA was fragmented and reverse-transcribed into cDNA. Following second-strand synthesis with dUTP, end repair, and adapter ligation, the second cDNA strand was digested with UNG enzyme. The resulting libraries were amplified, pooled, and subjected to paired-end sequencing on an Illumina platform.

Gene expression was quantified (TPM) and aligned to the reference genome. Differential gene expression (*p* ≤ 0.05) was analyzed using edgeR. Enrichment analyses for GO terms and KEGG pathways were performed with GOatools and KOBAS, respectively (*p* ≤ 0.05), with key functional modules and regulatory pathways visualized.

#### Statistical analysis

All data were expressed as mean ± standard deviation and analyzed statistically using IBM SPSS Statistics 23. Each experiment was repeated three times.

## Acknowledgements

The authors would like to express their gratitude to Inner Mongolia Agricultural University for providing laboratory facilities. This work was supported by the Programs of Natural Science Foundation of Inner Mongolia (No. 2024MS03063), the Talent Cultivation Program for Outstanding Young Scientists and Technologists (No. SPY2025-5), and the First-Class Disciplines scientific research project (YLXKZX-NND-017).

